# Heterogeneity of single-cell mechanical responses to tumorigenic factors

**DOI:** 10.1101/078154

**Authors:** Aldo Leal-Egaña, Gaelle Letort, Jean-Louis Martiel, Andreas Christ, Timothée Vignaud, Caroline Roelants, Odile Filhol, Manuel Théry

## Abstract

Tumor development progresses through a complex path of biomechanical changes leading first to cell growth and contraction followed by cell de-adhesion, scattering and invasion. Tumorigenic factors may act specifically on one of these steps or have wider spectrum of actions, leading to a variety of effects and thus sometimes to apparent contradictory outcomes. Here we used micropatterned lines of collagen type-I/fibronectin on deformable surfaces to standardize cell behavior and to measure simultaneously cell size, speed of motion and the magnitude of the associated contractile forces at the level of a single cell. We analyzed and compared normal human breast cell line MCF10A in control conditions and in response to various tumorigenic factors. In all conditions, distinct populations of cells with a wide range of biomechanical properties were identified. Despite this heterogeneity, normal and transformed motile cells followed a common trend whereby size and contractile forces were negatively correlated with cell speed. Some tumorigenic factors, such as activation of ErbB2 or the loss of the beta subunit of casein kinase 2 (CK2), shifted the whole population towards a faster speed and lower contractility state. Treatment with transforming growth factor beta (TGF-β), induced some cells to adopt opposing behaviors such as extreme high contractility versus extreme low contractility. Thus, tumor transformation amplified the pre-existing population heterogeneity and led some cells to exhibit biomechanical properties that were more extreme than that observed with normal cells.

## Introduction

The malignant transformation of cells encompasses a complex sequence of events implicating many distinct pathways, thus making the process difficult to describe and to categorize. Throughout the development of a tumor, abnormal biochemical signaling, abnormal cell growth and changes in the mechanical properties within the tumor are closely connected and interdependent. For example, cell stiffness promotes cell growth (Klein et al., 2009), which increases tissue density and mechanical pressure, which in turn, triggers oncogenic pathways that further stimulate cell growth (Fernández-Sánchez et al., 2015). Moreover, the crosstalk between biochemical signaling and the mechanical environment evolve during tumor progression so that defined biochemical or mechanical signals may have distinct consequences depending on the stage of tumor development.

The mechanical properties of individual cells and the tumor tissue as a whole vary widely during tumor development, from the early cell-adhesion and cell-proliferation stages of tumor formation to the late cell-dissociation and cell-migration stages during tumor dissemination when cells have become more transformed (Plodinec et al., 2012; Weder et al., 2014). Biochemical signals promoting cell contractility may have distinct effects on tumor cells at the early stages of tumor development (e.g. proliferating cells) in comparison with tumor cells at later stages (e.g. metastatic/invasive cells; Aguilar-Cuenca et al., 2014; Fritsch et al., 2010). Therefore, the various crosstalks between signaling and mechanics at the distinct stages of tumor progression may underlie the heterogeneity of cell responses, and explain many conflicting results on the effects of oncogenic factors, cell mechanics and their interplay during tumor progression.

Although under specific conditions their roles have been clearly defined, the pleiotropic effects of the main oncogenic pathways are not universally observed in all tumors, preventing their use as prognostic markers. Conflicting results in carcinogenesis have been reported for thyroid hormones (Piekiełko-Witkowska, 2013), macrophage-colony-stimulating factor (Laoui et al., 2014), vascular endothelial growth factor (VEGF) (Zhan et al., 2009), epidermal growth factor (EGF) (Nicholson et al., 2001), and fibroblast growth factor-2 (FGF-2) (Korc and Friesel, 2009). Notably, transforming growth factor beta (TGF-β), which has been described as tumor suppressor and an oncogenic factor (Kubiczkova et al., 2012), appears to affect cell contractility in a manner dependent on substrate stiffness (Marinković et al., 2012). Moreover, TGF-β appears to increase the contractility of melanoma cells (Cantelli et al., 2015) but decrease the contractility of muscle cells (Mendias et al., 2012); and depending on substrate stiffness, TGF-β can increase or reduce the speed of cell migration (Wu et al., 2013), induce EMT or induce apoptosis (Leight et al., 2012).

The effects of cell contractility on tumor progression remain to be fully elucidated. Tissue stiffness is a hallmark of cancer (Paszek et al., 2005) and an enhancer of tumor growth (Fernández-Sánchez et al., 2015). However, the observation that cell contractility promotes tumor growth (Samuel et al., 2011) contrast with the recurrent observation of cancer cells being softer than normal cells (Cross et al., 2007, 2008; Efremov et al., 2014; Fritsch et al., 2010; Jonas et al., 2011; Li et al., 2008; Xu et al., 2012). Nevertheless the mechanical landscape within a tumor is complex reflected by the variations in stiffness within tumor tissue (Plodinec et al., 2012).

The relationships between the magnitude of contractile forces and metastatic potential of a given tumor cell may also vary between different types of tumor cells, the tumor-cell environment, and the types of measurements being made. Cell lines ranked with respect to increasing metastatic potential have been reported to produce traction forces that positively correlate (Jonas et al., 2011; Kraning-Rush et al., 2012) or negatively correlate with that potential (Indra et al., 2011; Munevar, 2001). However, this may reflect that metastatic potential is hard to quantify and cell migration speed or depth of invasion into collagen matrices are often considered as a good proxy for this potential. Nevertheless, migration speeds and invasive capacities have been positively correlated with stiffness and contractile forces in some studies (Indra et al., 2011; Mierke, 2013; Mierke et al., 2011a, 2011b; Rösel et al., 2008) but negatively correlated in others (Agus et al., 2013; Swaminathan et al., 2011). The interpretation of these relationships is further complicated by the fact that 2-dimensional speed and 3-dimensional speed may not necessarily be correlated (Indra et al., 2011). Moreover, differences in the relationship between cell contraction and migration speed may well arise from the non-linear relationship between cell speed and cell adhesion. There is an optimal adhesion strength promoting cell migration, and increases or decreases in adhesion strength from this optimum reduces cell speed (DiMilla et al., 1991; Gupton and Waterman-Storer, 2006). Again, the relationship between cell speed and cell adhesion can depend on the stage of the tumor (Weder et al., 2014) and the type of cell migration - mesenchymal or amoeboid (Friedl, 2004) - making it difficult to compare results generated in different experimental conditions.

The averaging of values in cell populations may also mask important cell heterogeneity that exists within a tumor (Lee et al., 2014; Mcgranahan and Swanton, 2015), such as differences in mechanical properties (Plodinec et al., 2012). Therefore, to better characterize the mechanical and biochemical relationships, it is necessary to measure all parameters on the same single cells and compare those values measured on individual cells or subgroups (Altschuler and Wu, 2010; Eberwine and Kim, 2015). Here we used a cell-culture system based on micropatterned cell-adhesion substrates as a platform to measure cell speed, cell size and cell contractile forces under controlled conditions, in order to compare the responses between individual normal cells and those submitted to tumorigenic factors.

## Results and Discussion

Cell-shape changes in motile cells induce large variability in biophysical measurements. In cell culture, micropatterned lines of cell-adhesion substrate have been shown to standardize cell behavior and better approximate to in vivo conditions than a planar coating of the cell-adhesion substrate (Doyle et al., 2009). Moreover, the linear (fibril-like) organization of extracellular matrix proteins have been shown to be characteristic of the metastatic niche (Artym et al., 2015; Gritsenko et al., 2012; Lu et al., 2012). So the cells were evaluated in polyacrylamide hydrogels with a Young modulus of 10 kPa, which corresponds to the stiffness of breast tissue (Rzymski et al., 2011), that were micro-patterned with 4 μm width lines made of collagen type I and fibronectin (Vignaud et al., 2014) (Figure 1A). The experimental system was designed to measure the size, speed and contractile forces of each individual cell (Bastounis et al., 2014) (Figure 1B). Contractile forces were measured using an automated approach based on an ImageJ plugin we have developed previously, whereby individual cells are tracked and the energy required for the traction force field is computed from the displacements of fluorescent beads embedded in the substrate (Martiel et al., 2015).

**Figure 1:**
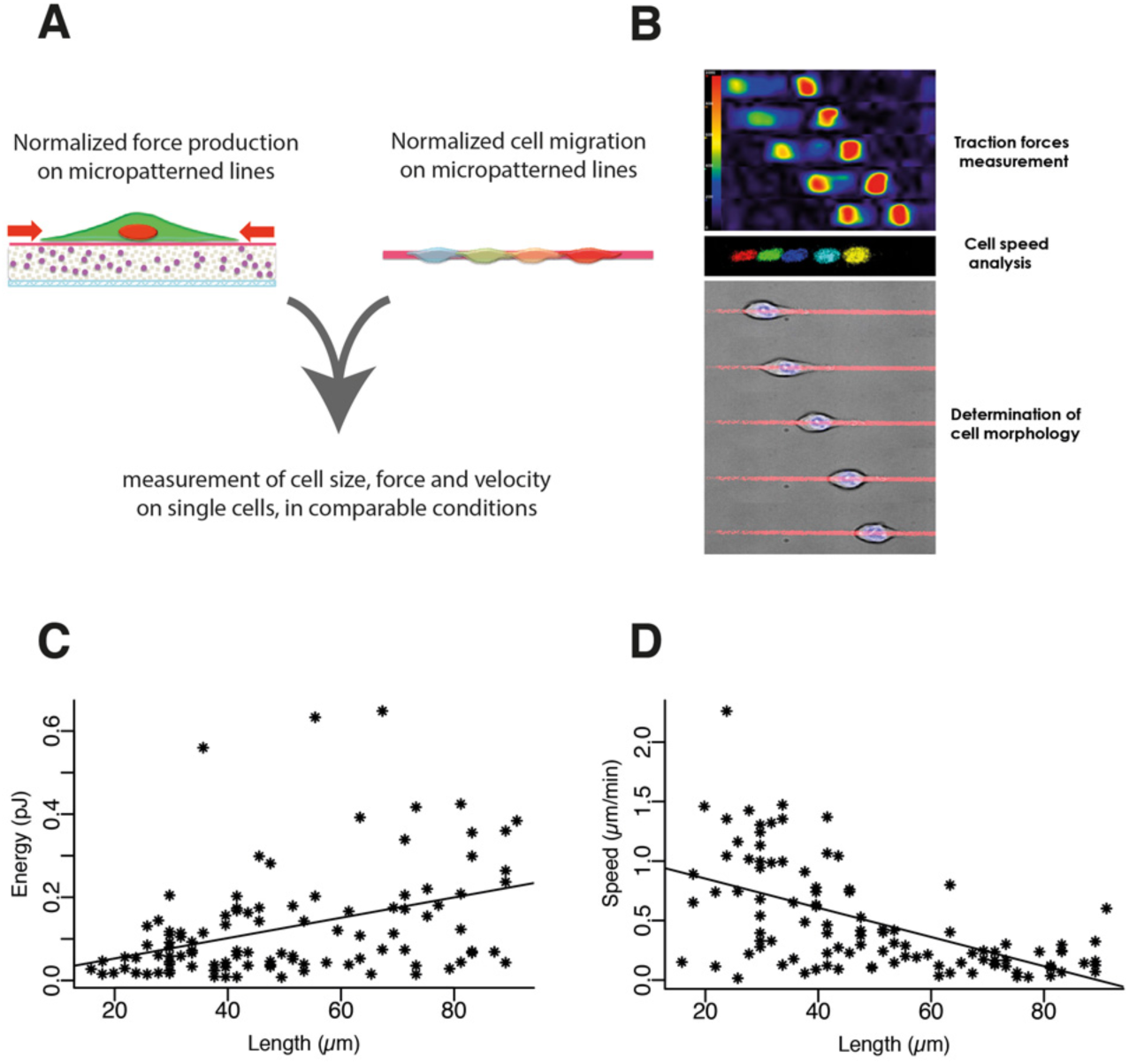
Methodological approach. A- Dynamic analyses of cell motion and traction forces exhibited by motile cells on micro-patterned polyacrylamide hydrogels. B- Examples of traction force field, nuclei displacements and cell motion. C- The graph shows contractile energy (measured by the energy required to deform the polyacrylamide hydrogel) plotted against cell length. The line describes the linear regression. Pearson correlation coefficient = 0.39, 95% confidence interval = 0.21 - 0.54 D- The graph shows cell speed plotted against cell length. The line describes the linear regression. Pearson correlation coefficient= −0.57, 95% confidence interval = −0.69 - −0.43

MCF10A cell line has been derived from non-transformed human mammary epithelium (Debnath et al., 2003). The cell line has been used in an established model of tumor transformation. In control conditions, the cells have the capacity to form acini-like structure in 3D gels. However, the cells are sensitive to oncogenic factors such as TGF-β and ErbB2 activation, which can force cells to grow in non-proliferative conditions (Debnath et al., 2003; Giunciuglio et al., 1995; Seton-Rogers et al., 2004; Xue et al., 2013). At baseline, the contractile energy (i.e. the mechanical energy used by the cell in substrate contraction) was positively correlated with the length of the cell (Figure 1C). By contrast, the speed of a motile cell was inversely correlated with the length of the cell (Figure 1D).

Furthermore, two distinct populations were identified based on the distribution of cell lengths (Figure 2A); one composed of *small* cells (shorter than 56 µm with an average length of 37 µm) and one composed of *large* cells (longer than 56 µm with an average length of 72 µm) (Figure 2A). Strikingly, cells from the small-cell subpopulation compared with cells from the large-cell subpopulation exhibited lower contractile forces and were faster (Figure 2B). Thus, the small-cell and large-cell subpopulations appeared to have distinct biophysical properties, which served as reference wild-type (WT) cell subpopulations in the study.

**Figure 2:**
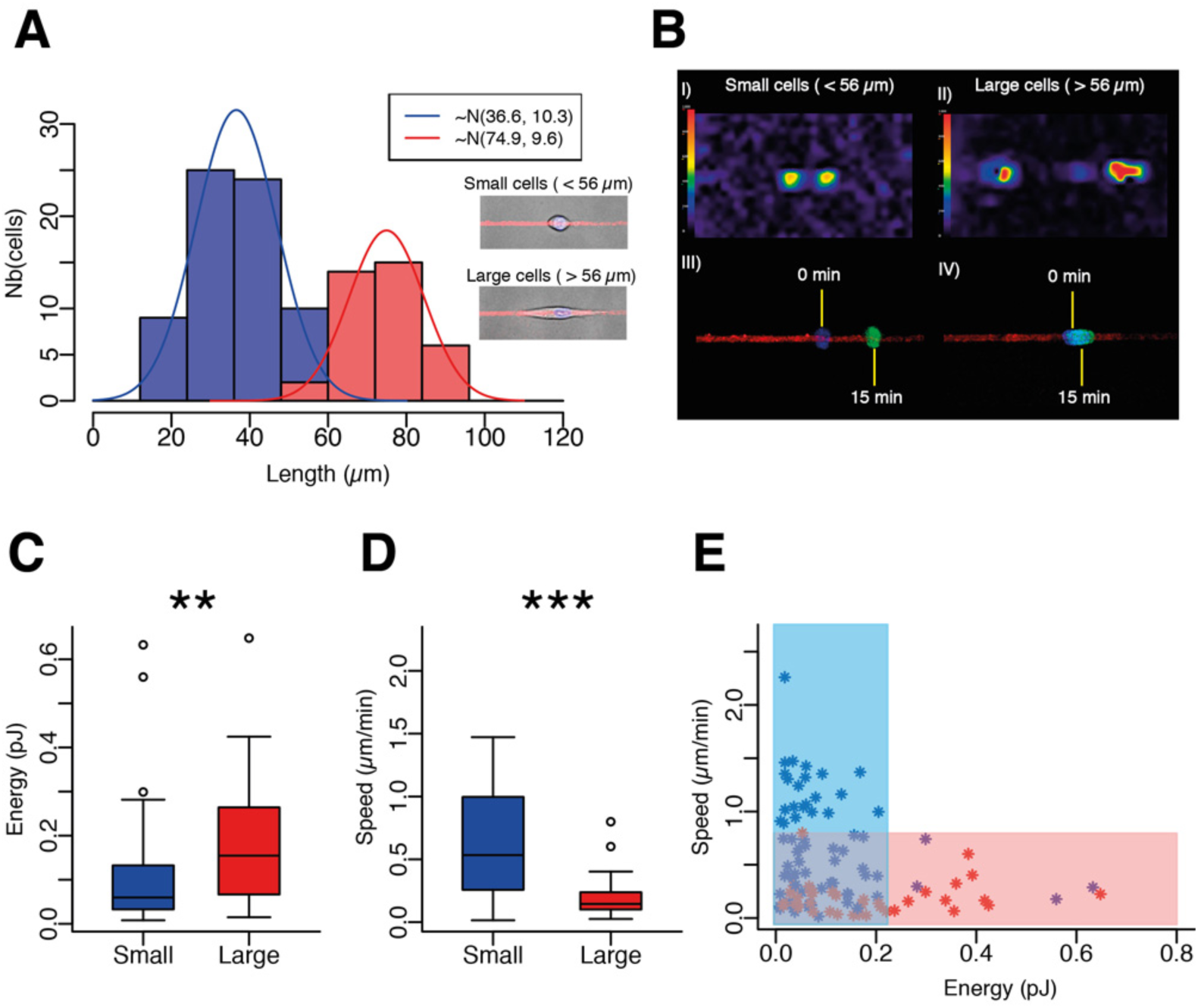
Analysis of cell sub-populations found within the MCF10A cell line (N=7 independent experiments, n=105 WT cells). A- Size distribution of WT cells. Two distinct cell subpopulations of cells were identified on the basis of size; *small* (blue) and *large* (red) with small cells being below 56 µm in length and large cells being greater than 56 µm in length. B- Representative examples of traction-force fields and corresponding nuclei displacements for small and large cells. C- Contractile energy of small versus large cells. D- Speed of small versus large cells. E- The graph shows cell speed versus contractile energy and the colored regions distinguish the low-contraction forces and high-speed phenotype (blue) from the high-contraction-forces and low-speed phenotype (red).

We then investigated the biophysical properties of a tumorigenic MCF10A cell line in which ErbB2 was constitutively active (Levental et al., 2009; Muthuswamy et al., 2001). These cells are characterized by an overexpression of the ErbB2 receptor tyrosine kinase, a receptor which has been identified as an activator of proliferation and invasiveness in breast cancer cells (Brix et al., 2014). In contrast to WT cells, the majority of ErbB2 cells were round and small with a peak length of 30 μm (Figure 3A), and only a minority of ErbB2 cells were large, with an average length of 85 μm (Figure 3B). The average size of ErbB2 cells in the small-cell subpopulation was also smaller than that in the small-cell subpopulation of WT cells. These ErbB2 cells were also faster and displayed less contractile forces than WT cells (Figure 3C,E). Interestingly the presence of small and low adhesive cells within surrounding tissues, has been described as symptomatic of metastasis and associated to poor prognosis in urinary bladder carcinomas (Cheng et al., 2004). The average size of ErbB2 cells in the large-cell subpopulation was similar to that in the large-cell subpopulation of WT cells (Figure 3D, F). Therefore the trend relating small cell size, high speed and low contractile forces observed in WT cells was apparent in ErbB2 cells, with an accentuation toward smaller and faster motile cells.

**Figure 3:**
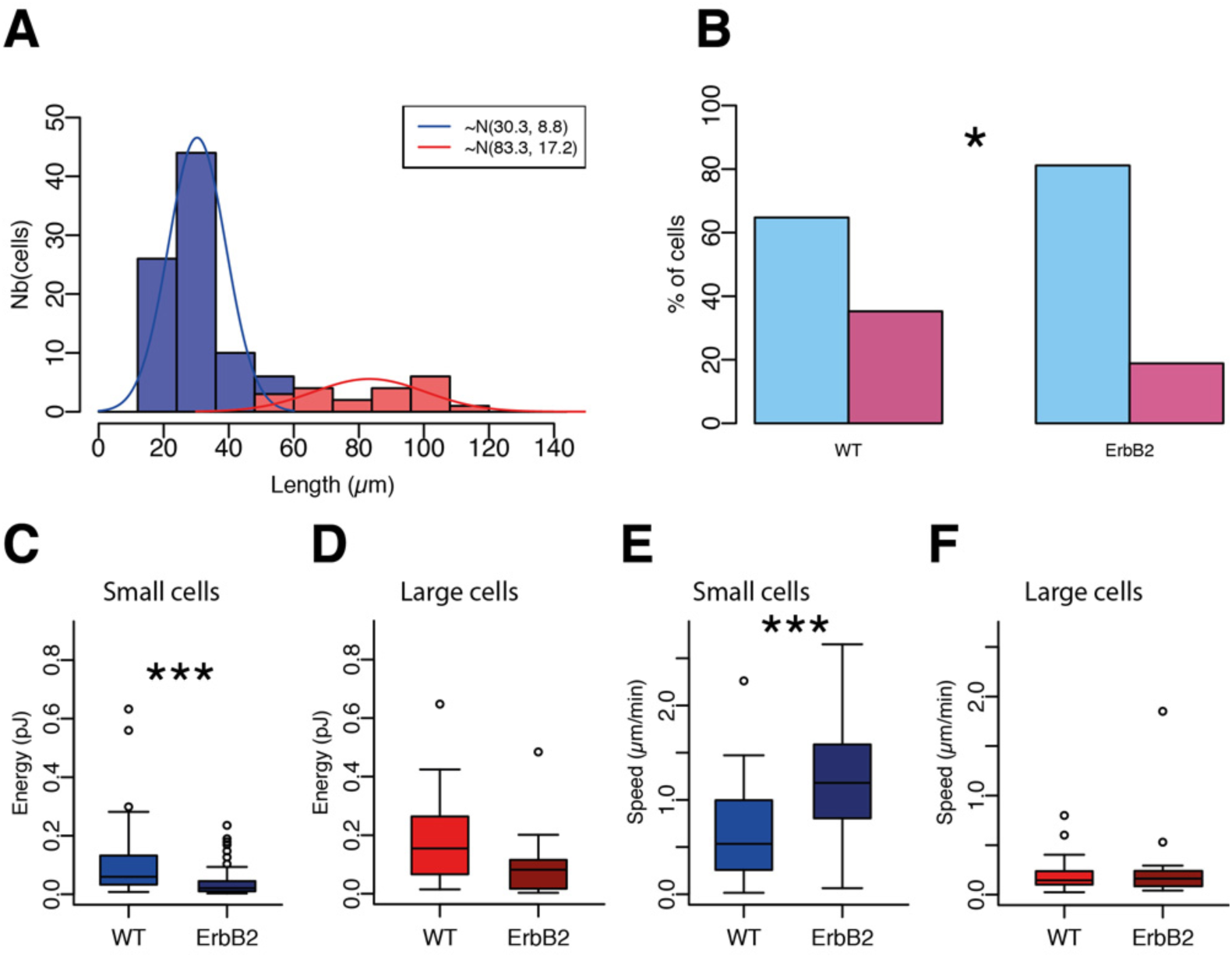
Analysis of morphology, contractility and migration speed of ErbB2 cells (N=3 independent experiments, n=106 cells). A- Size distribution of ErbB2 cells. Small cells are shown in blue, large cells are shown in red. B- Statistical analysis of population distribution according to cell length. C- Contractile energy of small cells. D Contractile energy of large cells. E- Small-cell speed. F- Large cell speed.

We then investigated the tumorigenic MCF10A cell line in which the beta subunit of the casein kinase 2 is depleted (MCF10A ΔCK_2_β). These cells are anti-apoptotic, pro-survival, multi-drug resistant and display EMT-like features (Deshiere et al., 2013; Vilmont et al., 2015). However, a direct role of casein kinase 2 depletion on cell metastasis and tissue invasion has yet to be identified. The distribution of ΔCK_2_β cell sizes fitted into a single population with a peak size of 70 μm (Figure 4A) and resembled the elongated morphology of large WT cells. By contrast and in biomechanical terms, the ΔCK_2_β cells resembled the small WT cells rather than the large WT cells, i.e. relatively high speed and low contractile forces (Figures 4B and 4C).

**Figure 4:**
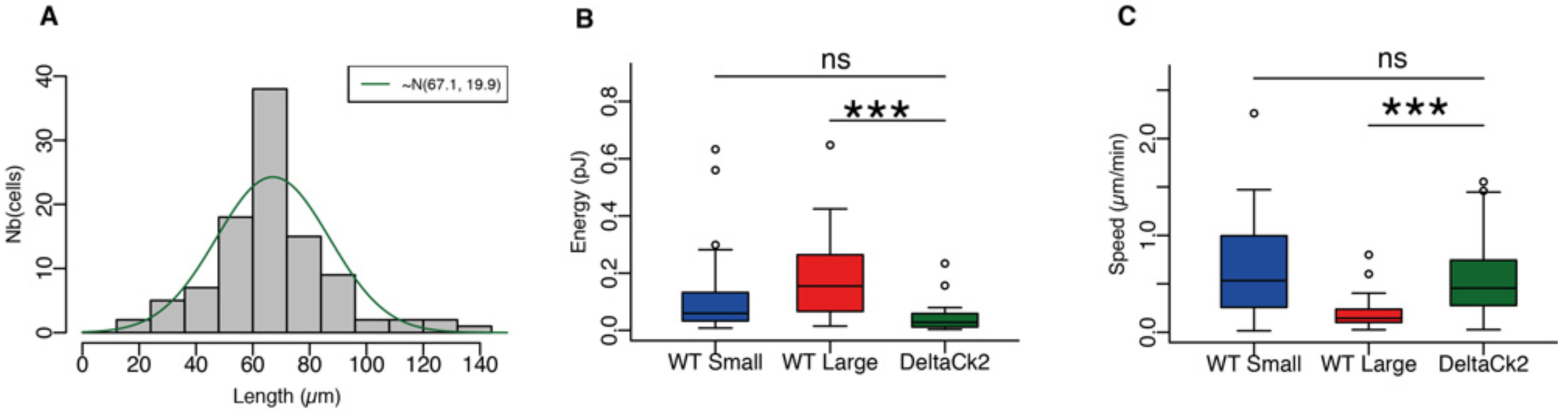
Speed and contractility of ΔCK_2_β cells (N=6 independent experiments, n=101 cells). A- Only a single population ΔCK2β cells were identified based on size distribution. B- Contractile energy of ΔCK2β cells compared with small and large WT cells. C- Speed of ΔCK2β cells in comparison with small and large WT cells.

We finally investigated the effect of TGF-β) on MCF10A cells by incubating the cells with 5 ng/mL TGF-β for 48 hours prior to start the experiment. In terms of morphology, TGF-β-treated cells resembled ErbB2 cells, with a majority subpopulation of small cells (peak length around 30 μm), and a minority subpopulation of large cells (average length of 90 μm). In biomechanical terms, the small TGF-β-treated cells also resembled ErbB2 cells and exhibited significantly higher speed and (marginally) lower contractile forces than small WT cells (Figure 5C). However, the large TGF-β-treated cells differed from ErbB2 cells in that the large TGF-β-treated cells exhibited significantly higher speed and higher contractile forces than large WT cells (Figure 5D). Notably, and unlike ErbB2 cells in comparison with WT cells, the differences in contractile forces between TGF-β-treated cells and WT cells was masked when the entire population of small and large cells were considered together (Figure 5E).

**Figure 5:**
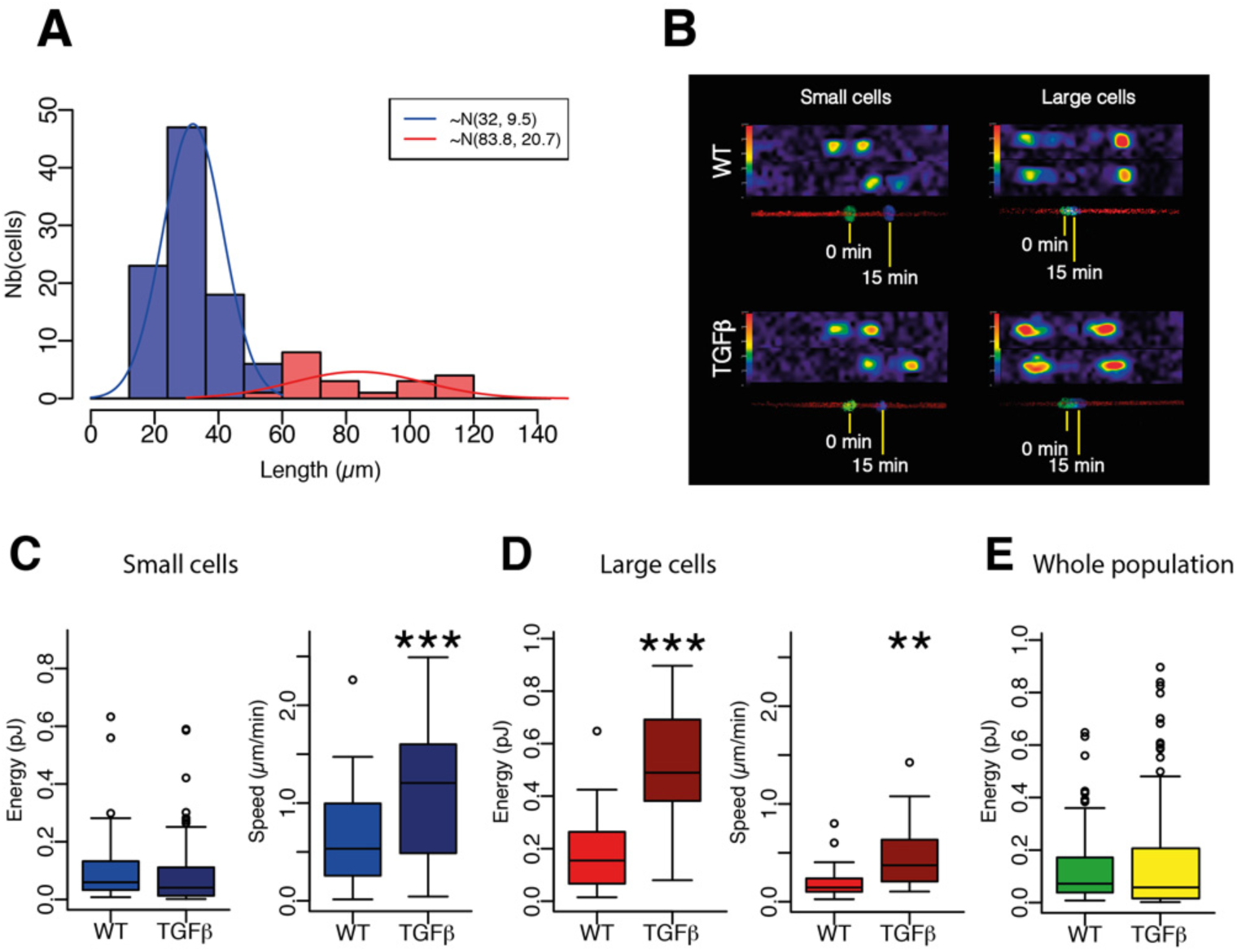
Speed and contractility of TGF-β treated cells (N=5 independent experiments, n=114 cells). A- Size distribution of TGF-β-treated cells. Small cells are shown in blue, large cells are shown in red. B- Representative examples of traction force fields and corresponding nuclei displacements. C- Contractile energy and migration speed of small cells. D- Contractile energy and migration speed of large cells E- Contractile energy of the entire populations considering together small and large cells.

Altogether our analyses showed that subpopulations in MCF10A-WT cells, ErbB2 cells and TGF-β-treated cells identified on the basis of differences in cell size also captured differences in contractile forces and speed. Given the relationship of ErbB2 and TGF-β to cancer, the differences in contractile forces and speed may reflect functional attributes that are relevant for transformed cells in vivo. In a further analysis, we superimposed the limits that defined the two subpopulations of WT cells on the cell speed versus contractile energy graphs (Figure 6). Hence in MCF10A-WT cells, the area defined by the blue rectangle set the limit for small cells (fast speed and low contractile forces), and the area defined by the red rectangle set the limit for large cells (low speed and high contractile forces).

Although significantly biased toward low contractile forces, MCF10A ΔCK_2_β cells appeared to be within the limits defined by the WT cells (Figure 6). By contrast, 40% of small ErbB2 cells and 50% of the small TGF-β cells were outside the limits defined by WT cells (blue arrows). Furthermore, almost 80% of large TGF-β-treated cells were outside the limits defined by WT cells (red arrows). Most of TGF-β-treated cells that exceeded the limit followed the expected trend in that the contractile forces opposed the migration speed. However, 15% (3/20) of large TGF-β-treated cells exhibited relatively high speed in addition to high contractile forces (area defined by dotted line), suggesting that the mechanism relating shape regulation, force production and cell motion was different in these cells compared with all other cells in this study.

**Figure 6:**
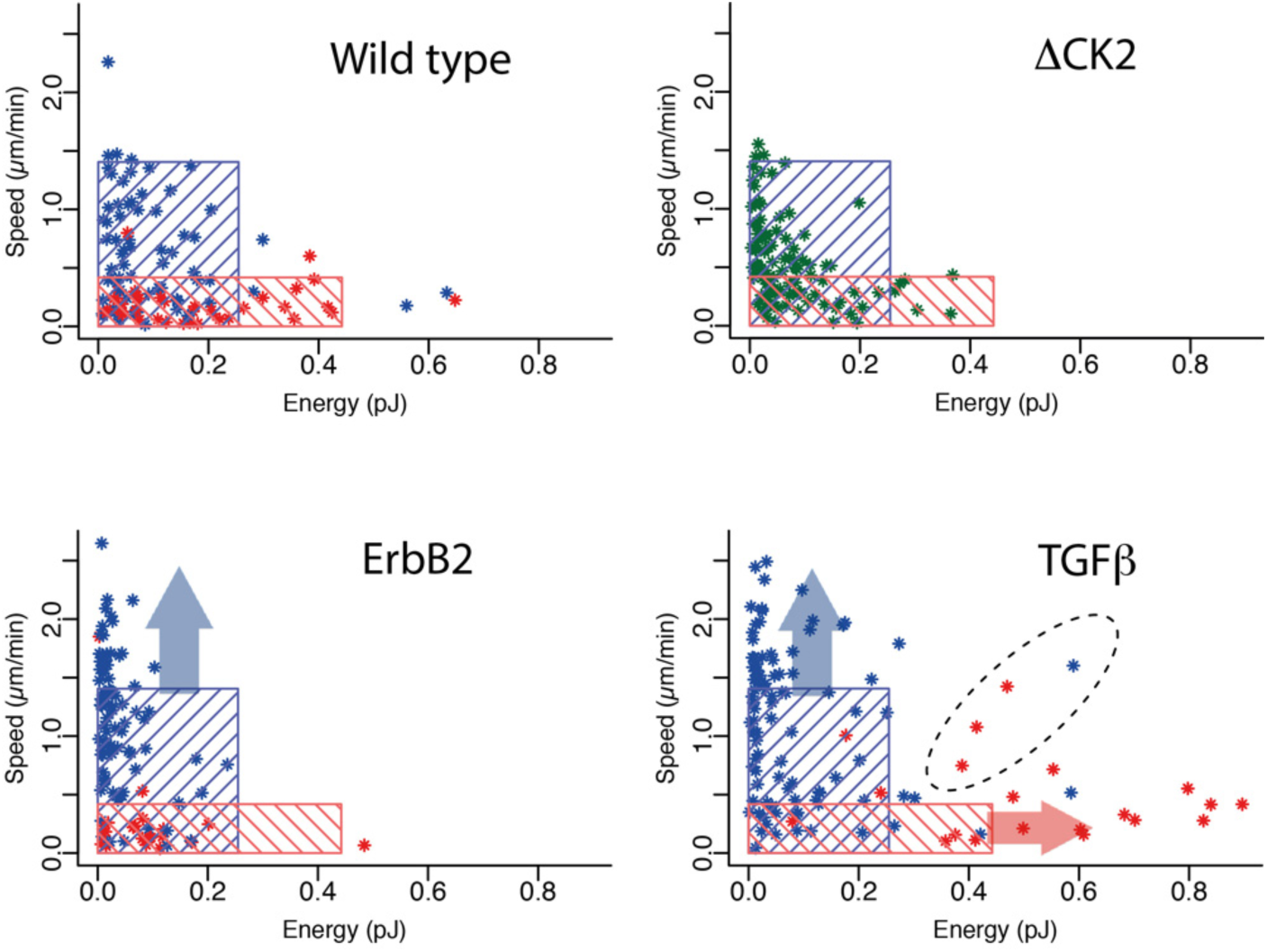
Heterogeneity of single cell response to tumorigenic factors. Graphs representing cell speed versus contractile energy. In the top-left graph, the blue rectangle includes 95% of the subpopulation of small WT cells, and the red zone includes 95% of the subpopulation of large WT cells. In the other three graphs representing the data from ΔCK2β cells, and TGF-β-treated cells, the same blue and red rectangles defined by the WT-cell subpopulations were superimposed. Blue arrows represent those small cells with relatively high speeds that are outside the limit defined by small WT cells. Red arrows represent those large cells with relatively high contractile forces, and which are outside the limit defined by large WT cells. The dotted line surrounds cells with relatively high speeds and contractile forces, and which in addition to be outside the limits defined by WT cells do not respect the global trend negatively relating cell speed and contractile energy.

## Conclusion

Our results identified two subpopulations of motile MCF10A cells based on size, which also had different mechanistic properties on collagen tracks. The small-cell subpopulation exhibited higher speeds and lower contractile forces, and we suggest that this subpopulation is likely to provide cells with a high metastatic and invasive potential. By contrast, the large-cell subpopulation exhibited lower speeds and higher contractile forces, and we suggest that this subpopulation is likely to provide cells which proliferate and support tumor growth and stiffness. Both normal and transformed cells followed a common trend whereby size and contractile forces were negatively correlated with cell speed. The tumorigenic factors, ErbB2 or the depletion of CK2, shifted the whole population of cells towards a faster speed and lower contractility state with some cells outside the high-speed boundary set by normal cells. Therefore the balance within a tumor between cell dissemination versus cell growth may reflect this overall shift in biomechanical properties (low to high speeds and high to low contractile forces). The response of MCF10A cells to TGF-β revealed that there may be further complexity. TGF-β induced some cells outside either or both the high-speed and high contractility boundaries set by normal cells, and included cells adopting opposing behaviors such as extreme high contractility versus extreme low contractility, or behaviors outside the general trend such as high speed and high contractility. Such complex outcomes, due to population heterogeneity and distinct effects on specific sub-populations, may account for some of the conflicting conclusions that have been reported so far. More importantly, our results highlight the absolute necessity of considering the heterogeneity of individual tumor-cell biomechanical profiles when characterizing oncogenic factors.

## Materials and Methods

### Experimental system

22×22 mm patterned polyacrylamide hydrogels were made according to methods previously published (Vignaud et al., 2014). Briefly, hydrogels were placed on 20×20 mm silanized coverslips, and patterned with 4.0 μm width / 1000 μm length lines made of collagen type-I (Sigma-Aldrich, USA) and fibronectin (Sigma-Aldrich, USA).

### Cells

Michigan Cancer foundation (MCF10A) human mammary gland cells (ATCC #CRL-10317) ^were maintained at 37°C at 5% CO_2_ in Lonza Mammary Epithelial Growth Medium (Lonza MEGM^ bullet kit #CC3150 without Gentamycin) in the presence of 100 ng/ml of cholera toxin (Sigma #C-8052) and 0.5% antibiotic-antimycotic (Life Technologies #15240062). Measurements of contractile energy were carried out using MCF10A wild type cells, MCF10A-ErbB2 cells, MCF10A-ΔCK2β cells and MCF10A wild type cells after 48 hours incubation with TGF-β (5 ng/mL, Sigma-Aldrich, USA). Prior to experimental set-up, cells were maintained overnight in an incubator (5% CO_2_, 95% humidity) then assessed in terms of speed, morphology and contractile energy.

### Analysis of hydrogel stiffness

The elasticity of the polyacrylamide (PAA) hydrogels was determined by atomic force microscopy (AFM) using established procedures (Frey et al., 2007). Briefly, cantilevers (PT-GS with a borosilicate sphere tip of radius 2.5 micrometers and nominal spring constant 0.06 N/m, Novascan Technologies, USA) were mounted on a Bioscope Catalyst AFM (Bruker) and calibrated using the thermal noise method. Force indentation curves were acquired for forces of 1 to 10 nN and fitted with a Hertz model to yield elastic moduli.

### Measurement of cell length, contractile energy and speed

Cell lengths were extracted from the traction force profiles by detecting the positions of force application sites. Contractile energy was expressed as the energy required to deform the 10 kPa polyacrylamide hydrogel (i.e. the traction force multiply by the gel deformation) and the measurements were carried out according to method previous published using the open source program ImageJ (Martiel et al., 2015). Briefly, bead displacements were determined using Particle Image Velocimetry (PIV) method with a window size of 3.96 µm. The corresponding contractile-energy was estimated with the FTTC method. Projection of the exerted forces along the patterned line were then calculated and analyzed automatically in Python. Cell speed was determined by identifying the position of the cell nuclei (DAPI stained) in consecutive pictures (15 minutes apart) with ImageJ.

### Image acquisition

Images were obtained using a spinning-disk confocal microscope (Nikon) with a 40X objective. Cells were imaged every 15 minutes for a period of at least 2 hours to control that cells were in a steady-state, but image analyses for speed and contractile energy measurements were performed on the first two images.

### Statistical analysis

#### Statistical analyses were performed using the statistical software R

Cell length distributions and subpopulation were defined from WT MCF10A cell. To distinguish the presence of subpopulation, we tested the normality of length distributions for 1, 2 and 3 subpopulations. In each case, the separation of cell lengths in sub-population clusters was done with a K-means algorithm. Each subpopulation was then tested for normality with Shapiro-Wilk normality test. For a single cluster or 3 clusters, the distributions failed the normality test (Shapiro-Wilk normality test, p-value < 0.0002 and p-value < 0.04 respectively), whereas 2-means clustering generated two normal subpopulations (p-value > 0.1). Moreover, this clustering defined the threshold length (56 µm) separating the small-sized cells from large-sized cells, and was the median value between the longest small cell and the smallest large cell.

The comparison of two populations of cells based on the frequencies of cell-size phenotypes within these populations (Figure 3B) was carried out using Fisher’s exact test. Results of this test were represented on the graphs with the following p-value thresholds: p-value >0.01 (ns), <0.01 (*), <0.001 (**), < 0.0001 (***).

The comparisons of populations of cells based on contractile activities or based on speeds (i.e. between small and large WT cells, and between WT and other cell lines) were performed using the Mann-Whitney test. Distributions were represented in a boxplot graph and results of this test were represented on the graphs with the following p-value thresholds: p-value >0.01 (°), <0.01 (*), <0.001 (**), < 0.0001 (***).

The rectangular areas on the cell speed versus contractile energy graphs were determined using 95 percentiles (threshold percentile values varied between 75 and 99 with little effect on the results) of speed and contractile-energy data obtained from the WT cell subpopulations (small and large, respectively). Fisher’s exact test was used to compare the number of outlying cells (out of WT rectangles domains, p-value >0.05 (°), <0.05 (*), <0.01 (**), < 0.005 (***)).

## Acknowledgments

This work was supported by the Institut National du Cancer (INCA PLBIO2011 to OFC and MT).

